# Characterizing the Genome of the Bumblebee *Bombus haematurus* – A Range-Expanding Pollinator

**DOI:** 10.1101/2025.08.29.673031

**Authors:** Saam Hasan, Jonathan S. Ellis, Paolo Biella, Andrew F. G. Bourke, Wilfried Haerty, J. Vanessa Huml, Mairi E. Knight

## Abstract

Over the past two decades, the blood-tailed bumblebee *Bombus haematurus* has expanded its range from Western Asia into Central Europe. It is among several members of the *Pyrobombus* subgenus of bumblebees to have undergone recent population expansions. A number of studies have investigated these other species for genomic signatures that might be associated with expansions. Currently there are no published studies describing whole genome resources for *B. haematurus* and no whole genome data from regions near the edge of the expanding front. Here, we sequenced samples of *B. haematurus* from part of its recently expanded range in Northern Italy to generate the first whole genome data for this species. The genome size of *B. haematurus* is in a similar range to that reported for closely related *Pyrobombus* species (∼300 mega base pairs), with a predicted ∼ 13,500 genes. At the location sampled, there was a comparatively low nucleotide diversity, low inbreeding evidenced through a negative inbreeding coefficient, *F*_IS_ (-0.14), and low percentage of genome covered by long runs of homozygosity, LROH (∼0.03%), compatible with a recent history of population expansion. Using nucleotide diversity and Tajima’s D, we also detected evidence of positive and balancing selection in developmental biology and signalling genes. Comparison with other members of the *Bombus* genus and *Pyrobombus* subgenus revealed similar biological processes to be associated with signatures of selection across the genus. Our comparison also identified similar trends of *F*_IS_ and LROH between *B. haematurus* and other *Pyrobombus* species.

## 1. Introduction

Over the past 25 years *Bombus haematurus* has expanded its range from the Middle East and south-east Europe (Reinig, 1974, Skhirtladze, 1988) to much of central Europe (Biella *et al*., 2021; Jeniœ *et al*., 2010; Sárospataki *et al*., 2005; Šima and Smetana, 2009; Straka, 2015). It is among several members of the subgenus *Pyrobombus* that have undergone recent population expansions (Looney *et al*., 2019; Richardson *et* al., 2018, Sheffield and Palmier, 2023), including its closest phylogenetic relative, *Bombus hypnorum* (Crowther *et al*., 2019; Huml *et al* 2021). This subgenus has been highlighted as a particularly interesting anomaly within the *Bombus* genus as many species within it are either maintaining or increasing in abundance and distribution in contrast to most other subgenus where the reverse trends are recorded (Arbetman *et al* 2017).

There is evidence from a wide range of taxa (e.g. plants, insects and birds) that species range expansions are associated with genomic changes detectable at population levels (e.g. Bragg *et al*., 2015, Keller *et al*., 2010, Ma *et al*., 2024, summarized by Hallatschek and Nelson, 2009, Hallatschek and Nelson, 2010). As a group, bumblebees have been comparatively extensively studied from a genomic perspective (e.g. Brock *et al*., 2021, Huml *et al*., 2021, Huml *et al*., 2023, Theodorou *et al*., 2018, Sang *et al*., 2024). These and other studies have reported evidence of associations between particular genomic regions and potential adaptation to various changing environmental conditions including temperature (Heraghty *et al*., 2022), urbanization (Theodorou *et al*., 2018) and elevation of habitat (Sun *et al*., 2021). These studies have provided important insights into the interactive dynamics between genetic variation and environmental change.

*B. haematurus* is under-studied in this context as there are no genome assemblies and only a small number of published whole genome resources with no accompanying publications. The only published study investigating the genomics of this species was limited to COI sequences (Biella and Galimberti, 2020). Given that the range-expansion of *B. haematurus* is associated with several kinds of macrohabitats and environmental changes (including climate) in the newly colonized regions (Biella *et al*., 2021, Šima, 2018), characterizing any genomic underpinnings would significantly expand our understanding of how this and other key pollinator species may be affected by, and are adapting to, common environmental risk factors such as climate change.

This study was conducted on samples of *B. haematurus* from northern Italy, i.e. part of its most recently colonized range, with the overall aims of generating new whole genome resources and carrying out preliminary genomic analyses comparing *B. haematurus* with other taxa within the genus and subgenus. The nature and quantity of these genomic metrics were then compared to those reported in other species of varying decline status to identify any associations unique to *B. haematurus* or shared with other expanding species. Our specific aims were to: (i) Use whole genome sequencing to generate a genome assembly for *B. haematurus*; (ii) Quantify key genetic metrics as indicators of diversity and inbreeding and compare with other relevant species; (iii) Detect whether regions show any initial strong signatures of selection and whether these share any commonality with other members of the *Pyrobombus* subgenus.

## 2. Materials and Methods

Ten *B. haematurus* individuals (all females) were collected from three locations in the Province of Gorizia in north-eastern Italy, during March 2023. Samples were initially frozen and stored at -80°C. DNA extraction and sequencing were carried out at the BGI Hong Kong Tech Solutions NGS. DNA concentration was evaluated using a Qubit Fluorometer (Thermo Fisher Scientific). The purity of the samples was measured using agarose gel electrophoresis (1% agarose gel run at 150V). Genomic DNA (1μg) was then fragmented using a Covaris ultrasonicator (Covaris). Fragment sizes ranging from 200 to 400 bp were selected using magnetic beads. This was followed by end repair, 3’ adenylation, and ligation of adapters to the adenylated 3’ ends. Amplification was carried out using PCR on the adapter ligated DNA fragments. The double stranded products were denatured using heat, followed by circularization through splint oligo sequence. The single stranded circular DNA was used to prepare the final library. After quality control, sequencing was carried out using the DNABseq G-400 platform (BGI Genomics Ltd.).

Fastq output files produced from the sequencer were quality checked using FastQC version 0.12.0 (Babraham Bioinformatics). Low quality reads were trimmed using Trimmomatic V0.32.in paired-end mode (for paired-end sequence data, Bolger *et al*., 2014). Trimmed fastq files for all 10 samples were then pooled to aid the assembly (as regions not sequenced in certain samples may have been sequenced in others). ABySS (Simpson *et al*., 2009), an assembler suitable for eukaryotic genomes (Hayes *et al*., 2024), was used for the assembly step. Successively increasing k-mer sizes from 60 onwards were used until no further improvement in assembly quality was obtained. The assembled genome was annotated using Liftoff, with all parameters set to default (Shumate and Salzberg, 2020). Liftoff requires an existing annotation for the same or a closely related species. For this, the *B. hypnorum* (phylogenetically closest to *B. haematurus*, Cameron *et al*., 2007) annotation (Crowley *et* al., 2023a, Harrison *et al*., 2023) was used.

The Burrows-Wheeler algorithm (BWA) (Li and Durbin, 2009) was used for genome alignment using the BWA-MEM algorithm. *B. haematurus* genomes were aligned to the *B. hypnorum* reference genome (iyBomhypn1.2), chosen in the absence of a *B. haematurus* reference genome for its phylogenetic closeness and good quality assembly and annotation (Crowley *et al*., 2023a). SNPs were called from the generated alignments using the Genome Analysis Toolkit (GATK) version 4.6.0.0 (Auwera *et al*., 2013). The most up-to-date GATK best practice workflow for genomic variant calling (Broad Institute, 2024) was followed with two exceptions: both base and variant quality recalibration were not possible due to a lack of the required pre-existing variation database for bumblebees. Called variants were further filtered to retain those with quality >20, minimum depth ≥10, maximum depth ≤100, and presence in ≥90% individuals.

All samples were then checked for relatedness using KING 2.3.2 (Manichaikul *et al*., 2010) and VCFtools 0.1.16 (Danecek *et al*., 2011), where a kinship coefficient <0 indicated no familial relatedness. If detected, only one member from sibling pairs was retained for calculation of nucleotide diversity (π) and Tajima’s D across 10 Kb windows as measures of selection. These two widely used methods were employed instead of alternative methods because they do not require information such as population history, haplotype phasing, etc. that were lacking in a single location study of a previously un-sequenced species. Indications of inbreeding were assessed by calculating the extent of long runs of homozygosity (LROH) and the inbreeding coefficient (*F*_IS_) using VCFtools and Hierfstat (de Meeûs and Goudet, 2007), respectively. Before calculating *F*_IS_, variants were pruned using Plink (Purcell *et al*., 2007) to remove sites in linkage disequilibrium (LD) and ensure the independence of analysed loci. Confidence interval of *F*_IS_ was assessed by conducting bootstrapping for 1000 cycles. LROH analysis followed the parameters implemented by Sang *et al*., (2024). For nucleotide diversity (π) and Tajima’s D, windows with low values indicate positive selection and high values indicate balancing selection (Korneliussen *et al*., 2013, Pollard *et al*., 2006, Schrider, 2020). These outlier windows were identified using z-scores if the values followed a normal distribution or using the bottom and top 5% ranking if non-normal. Windows were further filtered to retain those containing ≥5 SNPs. To test if particular biological functions were enriched within windows exhibiting evidence of selection, gene ontology analysis was carried out for all genes present in those windows, against a background of all genes with called sites (variant + non-variant). DAVID (Huang *et al*., 2007, Sherman *et al*., 2022) and topGO (Alexa and Rahnenführer, 2024) were used for the ontology analysis. Additionally, over-representation of types of biological pathways within windows evidencing selection was tested using pathway data from Reactome (Orlic-Milacic *et al*., 2024). Using the Reactome database and hierarchical classification system, biological pathways associated with genes present in selection windows were grouped into 28 broad categories or top-level pathways.

Over-representation was tested using Fisher’s Exact Test and subsequent Bonferroni correction, comparing the selection windows against a background comprising all genes with called sites (the same as for the enrichment analysis).

## 3. Results

FastQC reported satisfactory sequencing quality for all samples (sequences and sample information available under BioProject accession PRJNA1110219). The total estimated genome length from all assembled reads (contigs) was 335 Mb and the estimated length from all contigs >1000 bp was 235 Mb (Table 1). The scaffold N50, the most commonly used metric for evaluating genome assembly quality (Jauhal and Newcomb, 2021), was 98,461 bp for kmer size=85 (best obtained value). Liftoff predicted 13,543 coding genes (protein plus RNA) from the assembled genome (Table 1), within the 13,000-21,000 range for coding genes reported from other *Pyrobombus* species (e.g. Heraghty *et* al., 2020, Crowley *et al*., 2023a).

**Table 1:** Key statistics for B. haematurus genome assembly. Contig – contiguous sequence, N50 – minimum contig length covering 50% of assembled genome, scaffold – best quality assembly that was generated. N’s – unknown bases.

| Assembly Statistic | Value |
| --- | --- |
| Total Genome Length (All Contigs) | 3.35E+08 |
| Total Genome Length (Contigs>1 Kb) | 2.35E+08 |
| Scaffold N50 | 98461 |
| Number of Scaffold Contigs | 13047 |
| GC Content (% of total bases) | 37.7 |
| N's Per 100 Kb | 66.43 |
| Number of Coding Genes | 13543 |

All individuals barring one (FR11-1A) had over 99% reads mapping to the *B. hypnorum* reference genome, with coverage ranging from >9× for sample FR11-1A to >30× for all other samples. The number of called variants after filtering for quality, depth, and presence in ≥90% individuals was 2,289,948. Both VCFtools and King returned negative kinship coefficients for every pair of individuals, indicating that the sampled bees were not related.

The average nucleotide diversity across the genome was 0.00025, 1 - 2 orders of magnitude lower than values reported from other bumblebees (Colgan *et al*., 2022, Larragy *et al*., 2023, Lozier, 2014, Theodorou *et al*., 2018). Values of nucleotide diversity (π) were not normally distributed, hence the bottom and top 5% ranked values were used to shortlist windows exhibiting positive and balancing selection respectively. This shortlisted 868 low-diversity and 995 high-diversity windows. None of the low-diversity windows passed the minimum 5 SNP filter and so they were not used for the ontology analysis. All high-diversity windows contained more than 5 SNPs. Tajima’s D values displayed a normal distribution, appropriate for applying z-scores. Based on z-score outliers, positive selection and balancing selection were evidenced in 278 and 616 windows, respectively. A total of 603 LROH of length 100 kb or longer were identified, encompassing 0.03% of the genome, The *F*_IS_ for the population, after bootstrapping, was -0.145 (confidence interval from -0.143 to -0.148). The ontology analysis identified 21 and 19 GO terms (biological processes) enriched within windows showing evidence for positive selection and balancing selection, respectively (Figure 1A, 1B).

**Figure 1.**
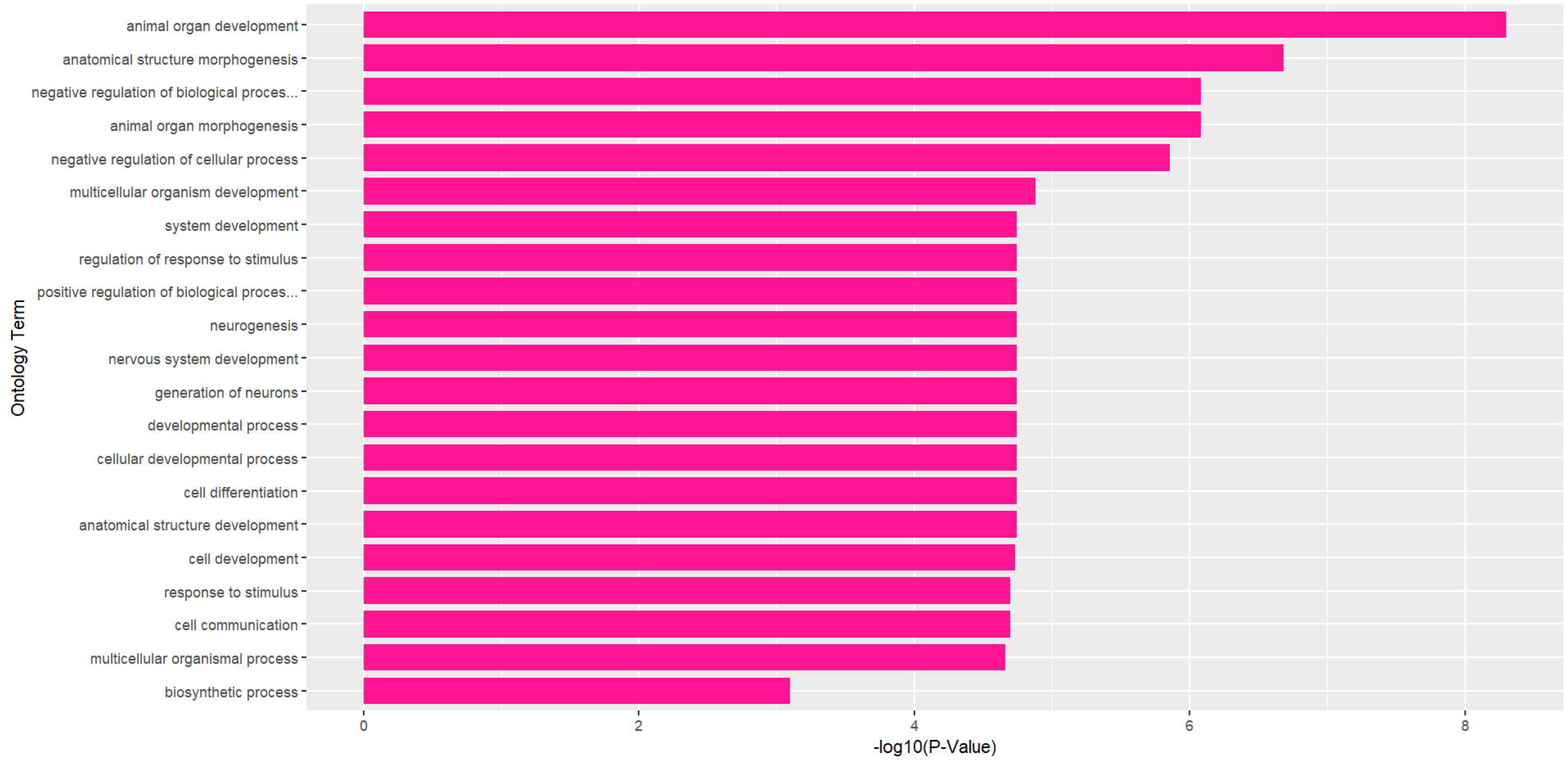
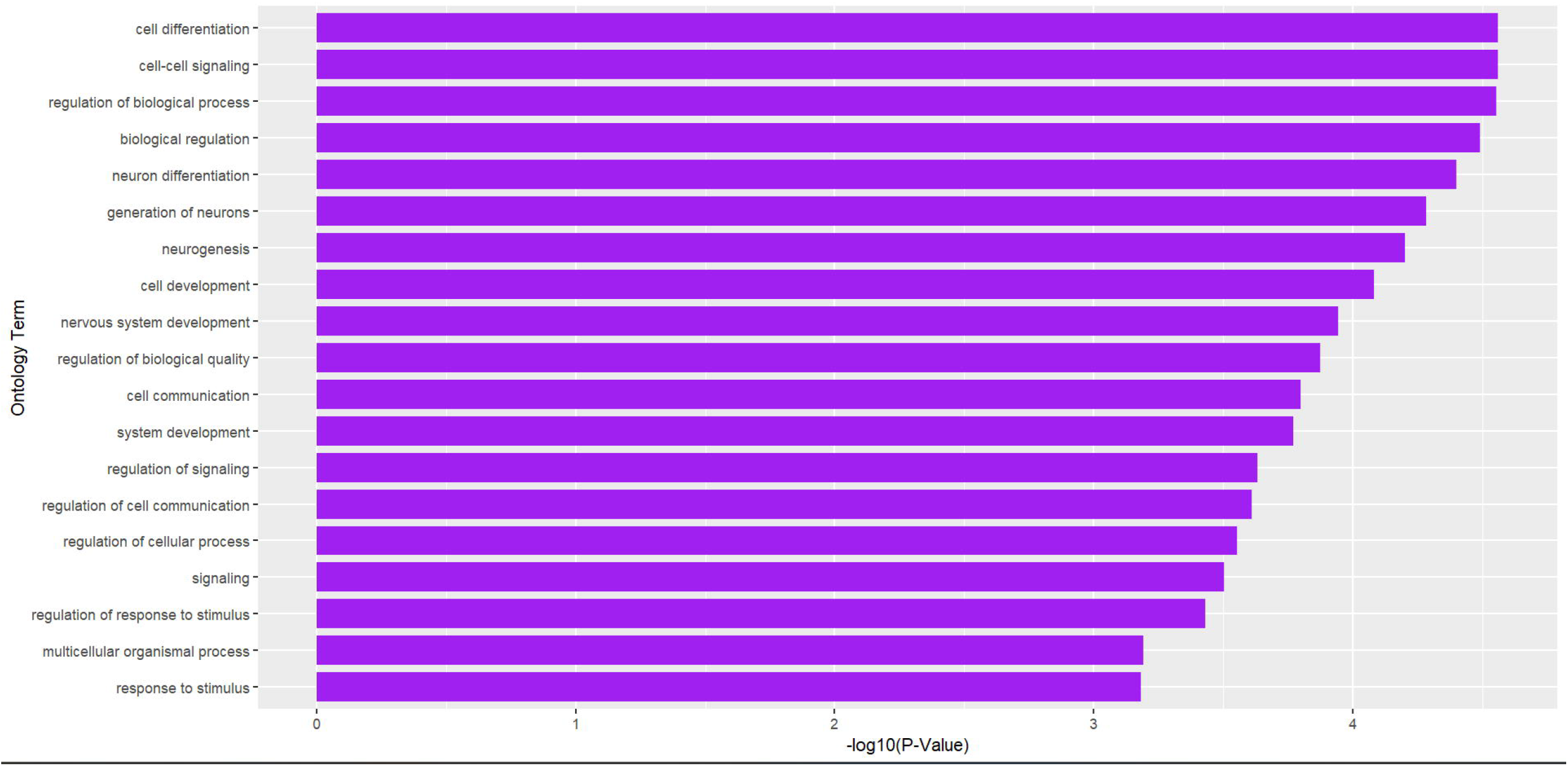
A and B: GO terms representing biological processes enriched at regions showing signatures of positive selection based on low Tajima’s D (1A). GO terms representing biological processes enriched at windows showing signatures of balancing selection (high Tajima’s D and high nucleotide diversity) (1B).

Twenty-eight pathway categories from Reactome were present in the regions of interest. From the Fisher’s Exact tests and subsequent Bonferroni correction, 8 categories for the positive selection windows and 9 categories in the balancing selection windows showed significant over-representation (Table 2).

**Table 2:**
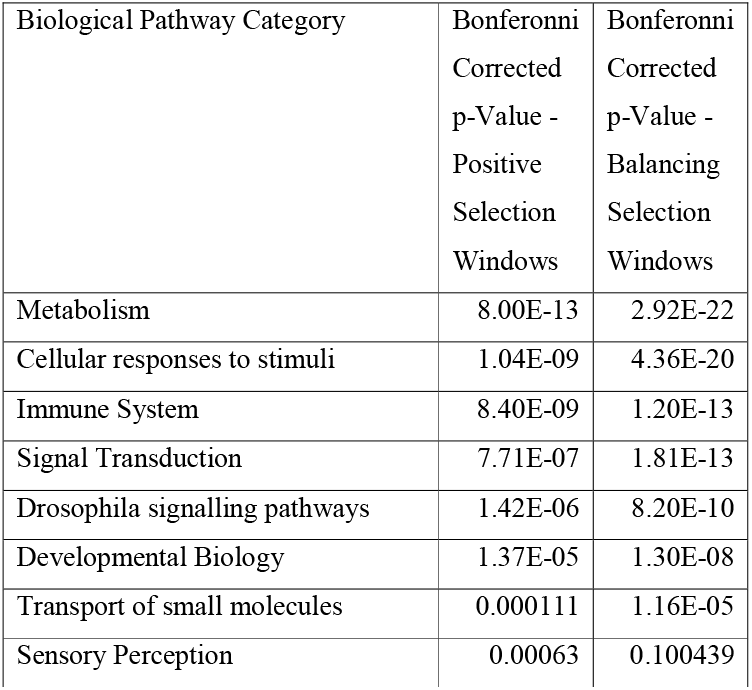
Over-representation of top-level biological pathways (biological pathway categories) within the windows exhibiting positive and balancing selection. Bonferonni-corrected p-value < 0.05 indicates significant over-representation.

| Biological Pathway Category | Bonferonni Corrected p-Value - Positive Selection Windows | Bonferonni Corrected p-Value - Balancing Selection Windows |
| --- | --- | --- |
| Metabolism | 8.00E-13 | 2.92E-22 |
| Cellular responses to stimuli | 1.04E-09 | 4.36E-20 |
| Immune System | 8.40E-09 | 1.20E-13 |
| Signal Transduction | 7.71E-07 | 1.81E-13 |
| Drosophila signalling pathways | 1.42E-06 | 8.20E-10 |
| Developmental Biology | 1.37E-05 | 1.30E-08 |
| Transport of small molecules | 0.000111 | 1.16E-05 |
| Sensory Perception | 0.00063 | 0.100439 |

## 4. Discussion

The genome size estimated for *B. haematurus* (Table 1) from the larger and consequently better assembled contigs (>1 kb), i.e. 235 Mb, was smaller than those reported from other closely related bumblebees (e.g. ∼297 Mb for *B. hypnorum*, Crowley *et al*., 2023a, 393 Mb for *B. terrestris*, Crowley *et al*., 2023c), although the size based on all contigs was closer to them. An underestimated genome size from large contigs could have stemmed from difficulty in filling gaps in the larger contiguous sequences with short-read data, but was nonetheless similar to sizes reported for other species using short-read technologies (e.g. *B. sylvicola* - 252 Mb, Christmas *et al*., 2021; *B. polaris* - 245.8 Mb, Sun *et al*., 2021). An underestimate could also be a consequence of the large number of repetitive regions known to be prevalent in bumblebee genomes (Sadd *et* al., 2015, Zhao *et al*., 2018), which complicate assembly by being incorrectly fused together into contiguous regions (Baptista *et al*., 2018). The number of predicted genes (∼13,500) was comparable to those reported for other bumblebee species using long-read data (Crowley *et* al., 2023b, Crowley *et al*., 2023c, Sun *et al*., 2021), although the closely related *B. hypnorum* has a considerably higher gene count (∼20,000; Crowley *et al*., 2023a). The lower observed gene count in *B. haematurus* could be due to the inability to assemble the contiguous sequences of a large number of genes from the short-read data. Additionally, although powerful for annotating genomes of species lacking reference genomes, due to its designed purpose of characterizing varying gene repertoires of populations of the same species, Liftoff may fail to annotate genes unique to *B. haematurus* or homologs/orthologs with high divergence rates (Shumate and Salzberg, 2021).

The negative inbreeding coefficients and low LROH detected in this study are compatible to what has been observed in expanding and common *Bombus* species, including other *Pyrobombus* species with similar demographic trends (stable/common/expanding), for example *B. hypnorum* (Huml *et al*, 2023) and *B. lapponicus* (Liu *et al*., 2023). In contrast, populations of *Bombus* species that are known to have experienced declines often exhibit positive inbreeding coefficients and/or high LROH occurrence (Charman *et* al., 2010, Kent *et* al., 2018, Mola *et* al., 2024, Sang *et al*., 2024).

The overall low nucleotide diversity was the most striking feature of the *B. haematurus* genome; >10 times lower than in other studied bumblebees (Colgan *et al*., 2022, Larragy *et al*., 2023, Lozier, 2014, Theodorou *et al*., 2018). Low nucleotide diversity is indicative of an excess of rare alleles (because nucleotide diversity is an average of pairwise differences, rare alleles contribute little, Akhunov *et al*., 2010), typical of recent population expansions (Booker and Keightley, 2018). A signature of low nucleotide diversity indicating an excess of rare alleles (Saavedra and Peña, 2005) but a negative *F*_IS_ value and lack of LROH (∼0.03% of genome) fits with a population experiencing a short-lived founder effect followed by expansion, as is recorded for *B. haematurus* (Biella *et al*., 2021, Rasmont *et al*., 2017); although future genomic studies of multiple populations will allow a more nuanced picture to emerge.

For both positive and balancing selection windows, the ontology analysis identified enrichment in genes associated with developmental functions. Selection on developmental biology genes strongly influences life-history traits and adaptation to ecological factors associated with the given niche of a species (Ellis and Del Giudice, 2019; Hendrikse *et al*., 2007). Developmental biology terms similar to those enriched at the regions of interest in this study have also been reported to be under directional selection in other bumblebees (for example Colgan *et al*., 2022).

The windows evidencing selection also showed overrepresentation of top-level pathways, suggesting they are rich in genes associated with these pathways. Among the over-represented pathways were some, e.g. immunity, sensory function, responses to stimuli, and signal transduction, that participate in responses to external changes and stressors such as pathogens, insecticides, climate adversity, etc. (Russel and Lightman, 2019). These processes have been associated with positive selection across the genus (Colgan *et al*., 2022, Huml *et al*., 2023, Jackson *et al*., 2020, Kent *et al*., 2018, Sun *et al*., 2021), as well as more widely (Alberola-Ila and Hernández-Hoyos, 2003, Diaz *et al*., 2018, Shultz and Sackton, 2019).

In conclusion, our data suggest that the *B. haematurus* genome is similar in size to those of its close phylogenetic relatives, with a gene count in a similar range to other bumblebees, though this is difficult to confirm from short-read data alone. There was no indication of inbreeding and the extent of LROH was low. We observed signatures of positive and balancing selection in developmental genes, providing early indication of significant adaptive changes in response to newly colonized habitats. Lastly, genes within the regions showing evidence of directional selection showed strong association, directly or indirectly, with pathways that respond to external stressors such as pathogens and insecticides. Future studies that utilize additional population genetics approaches on range-wide samples will further build on these conclusions.

## 8. Data Accessibility and Statement

The sequenced *B. haematurus* genomes are available on the NCBI SRA database under BioProject accession PRJNA1110219.

## 9. Competing Interests Statement

The authors declare no competing interests.

10. Acknowledgements

This study was funded by a University Research Studentship courtesy of the University of Plymouth. We would also like to acknowledge the Mohamed bin Zayed Species Conservation Fund, Project 232530334, for their support toward sample obtainment.

## 11. Author Contributions

The study was co-designed by all authors. SH oversaw the sequencing of samples, carried out the data analysis, and led the manuscript preparation. MK, JE, PB, AB, VH, WH edited and guided the MS. MK secured funding for the project. PB carried out sample collection.

